# Loss of *Acta2* in cardiac fibroblasts does not prevent the myofibroblast differentiation or affect the cardiac repair after myocardial infarction

**DOI:** 10.1101/2021.05.20.445043

**Authors:** Yuxia Li, Chaoyang Li, Qianglin Liu, Leshan Wang, Adam X. Bao, Jangwook P. Jung, Joseph Francis, Jeffery D. Molkentin, Xing Fu

## Abstract

In response to myocardial infarction (MI), quiescent cardiac fibroblasts differentiate into myofibroblasts mediating tissue repair in the infarcted area. One of the most widely accepted markers of myofibroblast differentiation is the expression of *Acta2* which encodes smooth muscle alpha-actin (SMαA) that is assembled into stress fibers. However, the requirement of *Acta2* in the myofibroblast differentiation of cardiac fibroblasts and its role in post-MI cardiac repair were still not known. To answer these questions, we generated a tamoxifen-inducible cardiac fibroblast-specific *Acta2* knockout mouse line. Surprisingly, mice that lacked *Acta2* in cardiac fibroblasts had a normal survival rate after MI. Moreover, *Acta2* deletion did not affect the function or overall histology of infarcted hearts. No difference was detected in the proliferation, migration, or contractility between WT cardiac fibroblasts and *Acta2*-null cardiac myofibroblasts. Additional analysis identified that *Acta2*-null cardiac myofibroblasts had a normal total filamentous actin level and total actin level. *Acta2* deletion caused a unique compensatory increase in the transcription level of *Actg2* and a possible increase in the protein abundance of cytoplasmic actin isoforms. In conclusion, the deletion of *Acta2* does not prevent the myofibroblast differentiation of cardiac fibroblasts or affect the post-MI cardiac repair, and the increase in the expression of non-SMαA actin isoforms and the functional redundancy between actin isoforms are likely able to compensate for the loss of *Acta2* in cardiac myofibroblasts.

## INTRODUCTION

Among cardiovascular diseases (CVDs) which are the leading causes of death in western countries (5), acute myocardial infarction (MI) is one of the most deadly forms due to its unpredictability, fast disease progression, and poor prognosis (1). Following MI, cardiomyocytes in the infarcted myocardium are permanently lost, which greatly compromises the integrity of the infarcted ventricle wall (2). Quiescent cardiac fibroblasts are quickly activated after MI, characterized by massive proliferation and myofibroblast differentiation (3), which is believed to be induced by mechanical stress and cytokine stimulation (4, 5). Myofibroblasts mediate the formation of infarct scar through expressing high levels of extracellular matrix (ECM) proteins and ECM remodeling enzymes (3, 6). In addition, myofibroblasts are also known for the presence of actin stress fibers mostly composed of smooth muscle alpha-actin (SMαA) encoded by *Acta2* (7, 8). We recently reported that the myofibroblast state of cardiac fibroblasts was largely limited to the first week after MI, after which myofibroblasts further differentiated into matrifibrocytes, a newly identified fibroblast differentiation state lacking the expression of myofibroblast marker genes (3). The transient myofibroblast state of cardiac fibroblasts suggests that myofibroblasts are especially important in the early stage of post-MI cardiac repair. Indeed, depletion of myofibroblasts after MI increased the risk of cardiac rupture during the first week after MI (6).

Due to the importance of ECM in post-MI tissue healing, the role of cardiac fibroblasts/myofibroblasts in ECM production and remodeling has received special attention (9-12). However, the function of SMαA, the hallmark of myofibroblasts, has been largely overlooked. A couple of early *in vitro* studies manipulating SMαA level or activity found that SMαA promoted fibroblast contraction but inhibited cell migration (13, 14). A later study showed that in myofibroblasts SMαA stress fibers play an important role in the formation of supermature focal adhesions which are important for the anchorage of myofibroblasts to the surrounding ECM (15, 16). Several previous studies using mice with global *Acta2* deletion to investigate the function of SMαA in myofibroblasts in different disease models suggest that SMαA stress fibers may regulate or affect myofibroblast proliferation, motility, contractility, and ECM remodeling (17-19). However, some discrepancies are present among the results reported by these studies, which suggests that the role and necessity of SMαA in myofibroblasts of different origins may vary significantly. In addition, the deletion of *Acta2* in other cell types, such as the vascular smooth muscle cell, may also affect the interpretation of the function of SMαA in studies using global *Acta2* knockout (KO) mice.

Here, we studied the function of SMαA in cardiac myofibroblasts and post-MI cardiac repair using a novel tamoxifen-inducible cardiac fibroblast-specific *Acta2* KO mouse line and cardiac fibroblast lineage tracing. It was found that *Acta2* deletion in cardiac fibroblasts did not significantly affect the post-MI survival or cardiac function of mice. In response to MI or TGFβ treatment, *Acta2*-null cardiac fibroblasts underwent normal myofibroblast differentiation, which was likely at least partially due to the compensatory effect of other actin isoforms.

### RESULT

### Cardiac fibroblast-specific deletion of *Acta2* does not affect post-MI survival, cardiac function, or remodeling in survived mice

To generate a mouse line with cardiac fibroblast-specific tamoxifen-inducible *Acta2* deletion, mice with *Acta2* exons 5-7 flanked by *loxP* sites (*Acta2*^*fl/fl*^) were crossed with Tcf21-MerCreMer (*Tcf21*^*MCM/+*^) mice that also carry a Cre-dependent eGFP construct in the Rosa26 locus (*R26*^*eGFP*^) (**Figure 1A**). In these mice (*Tcf21*^*MCM/+*^;*Acta2*^*fl/fl*^;*R26*^*eGFP*^), the deletion of *Acta2* exons 5-7 and expression of eGFP in *Tcf21*-lineage traced cardiac fibroblasts can be induced by tamoxifen treatment. *Tcf21*^*MCM/+*^;*Acta2*^*fl/fl*^;*R26*^*eGFP*^ and *Tcf21*^*MCM/+*^;*R26*^*eGFP*^ mice were treated with tamoxifen for 5 days to induce Cre activity starting at 6 weeks of age and then subjected to surgeries to induce MI at 9 weeks of age (**Figure 1B**). To maximize the efficiency of *Acta2* deletion and reduce the negative effect of prolonged continuous tamoxifen treatment on the viability of mice, another course of tamoxifen treatment was applied to mice from 1 day before MI to 5 days after MI. A slightly lower survival rate was seen in *Tcf21*^*MCM/+*^;*Acta2*^*fl/fl*^;*R26*^*eGFP*^ mice than in *Tcf21*^*MCM/+*^;*R26*^*eGFP*^ mice during the first week after MI, however, the difference was not statistically significant as determined by Chi-Square Test (*X*^*2*^ (1, n = 245) = 0.0688, *p* = 0.79) (**Figure 1C**). Mice of both groups rarely died after the first week after MI (data not shown), which is consistent with other research showing that most of the fatality happens during the first week after MI when the tissue remodeling is highly active (6, 12). Moreover, the cardiac function of the two groups was not significantly different from each other on both days 7 and 28 after MI (**Figures 1DE**). To study the potential impact of *Acta2* deletion on the post-MI cardiac repair and remodeling, heart samples were collected from *Tcf21*^*MCM/+*^;*Acta2*^*fl/fl*^;*R26*^*eGFP*^ and *Tcf21*^*MCM/+*^;*R26*^*eGFP*^ mice on days 7 and 28 after MI and subjected to immunohistochemical staining (IHC) for Col1a1 and trichrome staining (**Figures 2AB**). No difference in the histology and fibrotic response of the heart was observed between *Tcf21*^*MCM/+*^;*Acta2*^*fl/fl*^;*R26*^*eGFP*^ and *Tcf21*^*MCM/+*^;*R26*^*eGFP*^ mice (**Figures 2A-C**), suggesting relatively normal tissue remodeling in *Tcf21*^*MCM/+*^;*Acta2*^*fl/fl*^;*R26*^*eGFP*^ mice after MI. The unaffected thinning of the left ventricle free wall and dilation of the left ventricle in *Tcf21*^*MCM/+*^;*Acta2*^*fl/fl*^;*R26*^*eGFP*^ mice also suggest normal contractility of cardiac fibroblasts lacking *Acta2* (**Figures 2ABD**). The infiltration of *Tcf21*-lineage traced cardiac fibroblasts into the infarct scar was also not affected by the loss of *Acta2* (**Figure 2A**), suggesting that the motility of cardiac fibroblasts is not affected by the lack of SMαA stress fibers. Together, these results suggest that the loss of SMαA in cardiac myofibroblasts does not affect post-MI cardiac repair or tissue remodeling.

**Figure 1.**
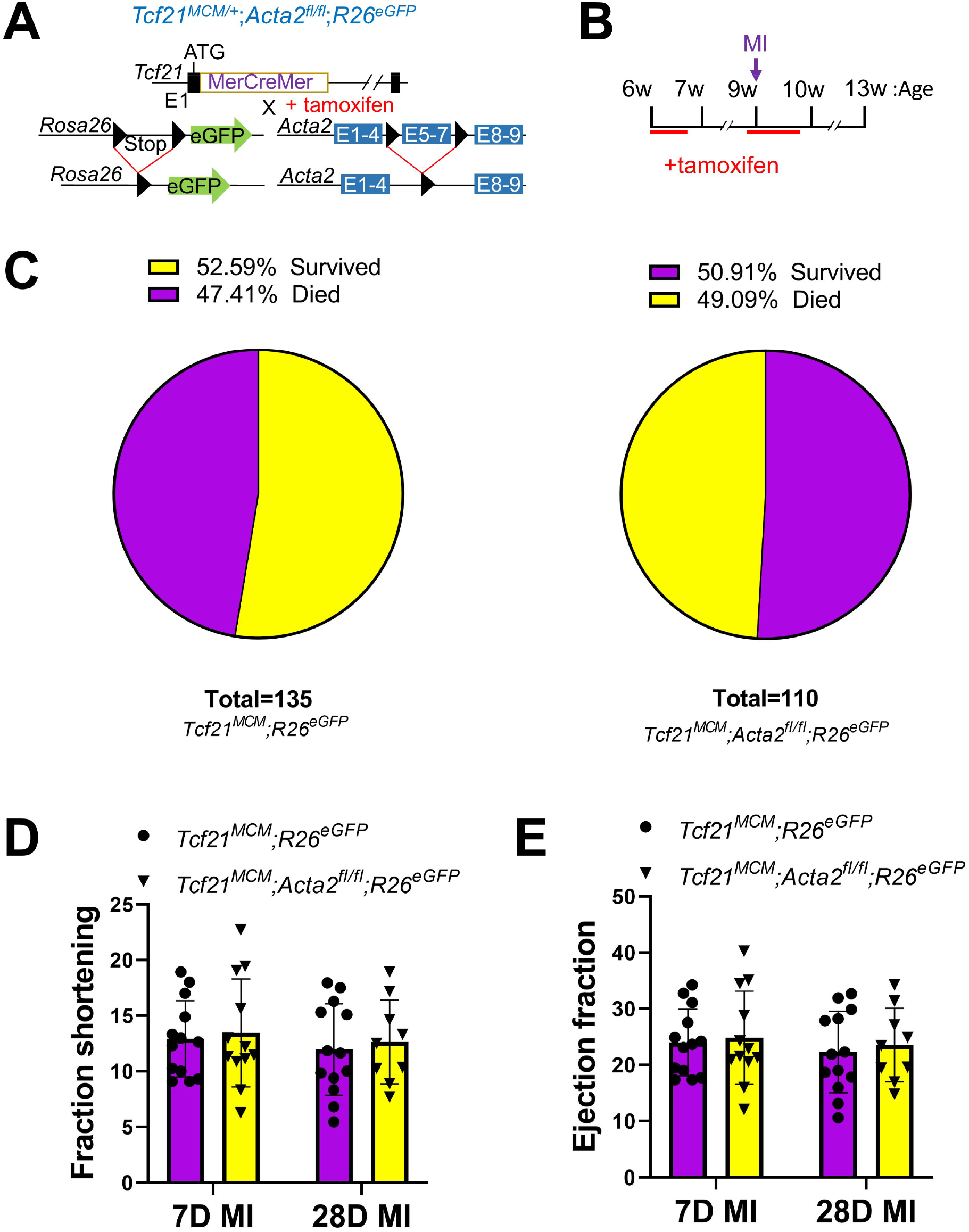
Cardiac fibroblast-specific deletion of *Acta2* does not affect the post-MI survival or cardiac function of mice. (A) Schematic of the generation of *Tcf21*^*MCM/+*^;*Acta2*^*fl/fl*^;*R26*^*eGFP*^ mice. In these mice, a tamoxifen-regulated MerCreMer cDNA cassette inserted into exon 1 (E1) enables the tamoxifen-induced deletion of the *loxP* site-flanked stop cassette upstream of *eGFP* inserted into the *R26* locus and the deletion of the *loxP* site-flanked exons 5-7 (E5-7) of *Acta2* only in cells expressing *Tcf21*. (B) Experimental scheme whereby *Tcf21*^*MCM/+*^;*R26*^*eGFP*^ and *Tcf21*^*MCM/+*^;*Acta2*^*fl/fl*^;*R26*^*eGFP*^ mice were given tamoxifen for 5 continuous days starting at 6 weeks of age and rested for 2 weeks before MI surgery. Mice were treated with tamoxifen again every day from one day before MI to 5 days. (C) The percentages of *Tcf21*^*MCM/+*^;*R26*^*eGFP*^ and *Tcf21*^*MCM/+*^;*Acta2*^*fl/fl*^;*R26*^*eGFP*^ mice survived or died within the first 7 days after MI. (D-E) Fraction shortening (D) and ejection fraction (E) of *Tcf21*^*MCM/+*^;*R26*^*eGFP*^ and *Tcf21*^*MCM/+*^;*Acta2*^*fl/fl*^;*R26*^*eGFP*^ mice at 7 and 28 days after MI.

**Figure 2.**
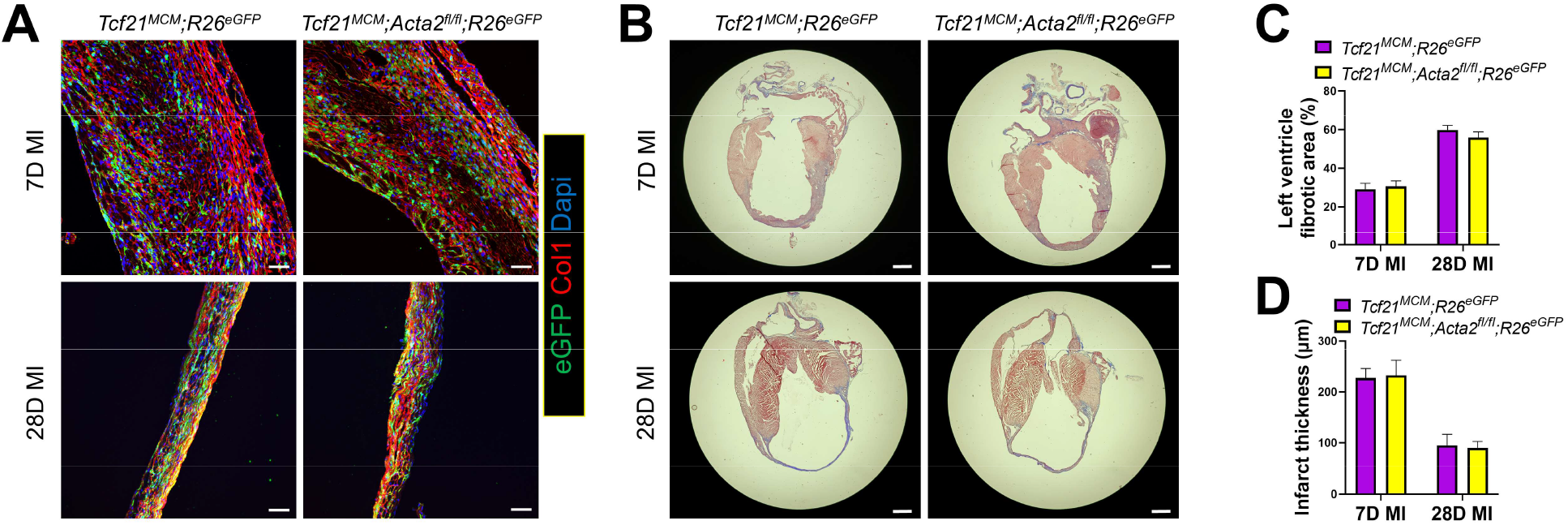
Cardiac fibroblast-specific *Acta2* KO mice have normal cardiac remodeling after MI. Heart samples were collected from tamoxifen-treated *Tcf21*^*MCM/+*^;*R26*^*eGFP*^ and *Tcf21*^*MCM/+*^;*Acta2*^*fl/fl*^;*R26*^*eGFP*^ mice at 7 and 28 days after MI. (A) Representative IHC images from 5 analyzed hearts per group showing the presence of eGFP^+^ *Tcf21* lineage-traced cardiac fibroblasts and the expression of type I collagen (Col1). Nuclei are shown with DAPI. Scale bar: 50 µm. (B) Representative trichrome staining images from 5 analyzed hearts per group showing the dilation of the left ventricle after MI and collagen deposition in the infarct area. Scale bar: 1 mm. (C-D) Percentage of the fibrotic area in the left ventricle (C) and average left ventricle thickness (D) determined by analysis of trichrome staining images using ImageJ. n=5.

### Cardiac fibroblast-specific deletion of *Acta2* does not affect the expansion of cardiac fibroblasts in the infarct after MI or their proliferation *in vitro*

It was reported that global *Acta2* KO led to increased renal fibroblast expansion (17). To study if specific deletion of *Acta2* in cardiac fibroblasts also affects their post-MI expansion, a 5-Ethynyl-2’-deoxyuridine (EdU)-based in vivo proliferation assay was performed. Using this method, we previously identified that the proliferation of cardiac fibroblasts after MI peaked at day 3 after MI and was largely diminished beyond day 7 post-MI (3). Thus, a single EdU injection was given to mice on days 3 or 7 after MI, followed by sample collection 4 hours after EdU injection. EdU and Ki67 staining showed that the post-MI proliferation of *Tcf21*-lineage traced cardiac fibroblasts was not different between *Tcf21*^*MCM/+*^;*Acta2*^*fl/fl*^;*R26*^*eGFP*^ and *Tcf21*^*MCM/+*^;*R26*^*eGFP*^ mice, despite the efficient cardiac fibroblast-specific *Acta2* deletion in *Tcf21*^*MCM/+*^;*Acta2*^*fl/fl*^;*R26*^*eGFP*^ mice indicated by the absence of SMαA expression in *Tcf21* lineage-traced cardiac fibroblasts (**Figure 3AB**). To verify these findings *in vitro*, cardiac fibroblasts were isolated from *Acta2*^*fl/fl*^ and WT mice, treated with adenoviruses expressing Cre (Adeno-Cre), and induced for myofibroblast differentiation by TGFβ treatment. No difference in the percentage of cells expressing Ki67 was observed between the 2 groups (**Figure 3C**). A proliferation assay was also performed by co-culturing TGFβ-and Adeno-Cre-treated cardiac fibroblasts isolated from *Acta2*^*fl/fl*^;*R26*^*eGFP*^ mice and *R26*^*tdTomato*^ mice (tdTomato is expressed in response to Adeno-Cre) mixed at a 1:1 ratio. The ratio remained the same after 48 hours of incubation, further suggesting that *Acta2* deletion does not affect the proliferation of cardiac fibroblasts (**Figure 3D**).

**Figure 3.**
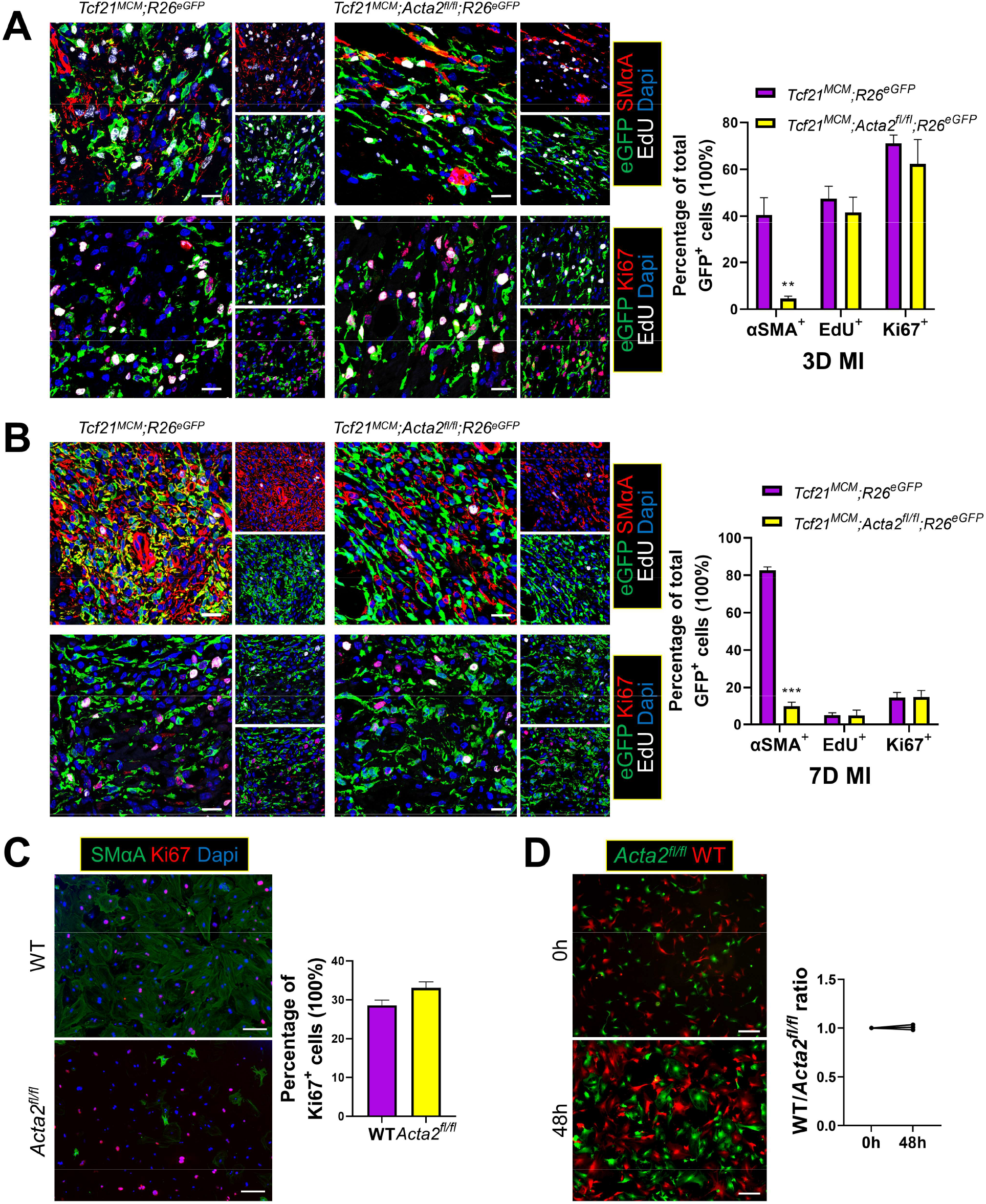
Cardiac fibroblast-specific deletion of *Acta2* does not alter cardiac fibroblast proliferation after MI. (A-B) Tamoxifen-treated *Tcf21*^*MCM/+*^;*R26*^*eGFP*^ and *Tcf21*^*MCM/+*^;*Acta2*^*fl/fl*^;*R26*^*eGFP*^ mice were subjected to MI. Heart samples were collected at 3 days and 7 days post-MI, 4 hours after a single EdU injection. IHC was performed to identify and quantify *Tcf21* lineage-traced (eGFP^+^) cardiac fibroblasts that are positive for SMαA, Ki67, or EdU in the infarct region at 3 days (A) and 7 days (B) after MI. n=3. Scale bar: 20 µm. (C) Cardiac fibroblasts isolated from WT and *Acta2*^*fl/fl*^ mice were transduced with Adeno-Cre and treated with TGFβ for 2 days. ICC was performed to identify the expression of SMαA and Ki67. Nuclei are shown with DAPI. n=3. Scale bar: 100 µm. (D) Cardiac fibroblasts isolated from *R26*^*tdTomato*^ (WT) and *Acta2*^*fl/fl*^*;R26*^*eGFP*^ (*Acta2*^*fl/fl*^) mice were transduced with Adeno-Cre and co-cultured at a 1:1 ratio. The ratio between WT and *Acta2*^*fl/fl*^ cardiac fibroblasts was calculated again after 48 hours of co-culture in the presence of TGFβ. n=3. Scale bar: 250 nm. \*\**P* < 0.01; \*\*\**P* < 0.0001.

### Cardiac myofibroblasts *lacking Acta2* have normal motility, contractility, and matrix stabilization ability

Besides the unaffected cardiac fibroblast proliferation, *in vivo* experiments using *Tcf21*^*MCM/+*^;*R26*^*eGFP*^ and *Tcf21*^*MCM/+*^;*Acta2*^*fl/fl*^;*R26*^*eGFP*^ mice also suggest that the deletion of *Acta2* in cardiac fibroblasts does not affect their motility and contractility after MI (**Figures 2A-D**). To verify these results in more controlled environments, the effects of *Acta2* deletion on the motility and contraction of cardiac fibroblasts were tested *in vitro*. For a direct comparison of motility, Adeno-Cre-treated *Acta2*^*fl/fl*^;*R26*^*eGFP*^ cardiac fibroblasts and *R26*^*tdTomato*^ cardiac fibroblasts co-cultured at 1:1 ratio were monitored for their migration toward the center of the well. No difference was observed between the 2 groups in the presence or absence of TGFβ (**Figure 4**). To better simulate the 3D matrix environment in the infarct, the same experiment was repeated with the addition of a layer of collagen gel cast on top of the cocultured cardiac fibroblasts. However, again no difference was observed (**Figure 4**). To test the effect of *Acta2* deletion on the contractility of cardiac fibroblasts, a gel contraction assay was performed using cardiac fibroblasts isolated from *Acta2*^*fl/fl*^ and WT mice and treated with Adeno-Cre and TGFβ, which showed no difference between the 2 groups (**Figure 5A**). It has been shown that cells can function as a crosslinker to strengthen the mechanical property of the matrix (20), a possible mechanism through which cardiac fibroblasts/myofibroblasts maintain the structural integrity of the infarct. Thus, we employed rheology to test the effect of cardiac fibroblasts on the stability of the collagen gel they reside in. In rheology, the storage modulus indicates the ability of the material to store deformation energy in an elastic manner while maintaining structural integrity and has been used to measure the strength of biomaterials (21-23). Hydrogels with a greater level of crosslinking have a higher storage modulus (24). Cardiac fibroblasts isolated from *Acta2*^*fl/fl*^ and WT mice were treated with Adeno-Cre and TGFβ, mixed with collagen gel, and culture for 24 hours to allow for the development of tension, followed by tests using a rheometer. The presence of cardiac fibroblasts increased the storage modulus of collagen gels (**Figure 5B**). However, no significant difference was observed in the storage modulus between collagen gels laden with WT myofibroblasts and those laden with *Acta2*-null myofibroblasts. This suggests that the loss of *Acta2* does not affect the matrix stabilization capacity of myofibroblasts, which is consistent with the similar survival rates between *Tcf21*^*MCM/+*^;*R26*^*eGFP*^ and *Tcf21*^*MCM/+*^;*Acta2*^*fl/fl*^;*R26*^*eGFP*^ mice.

**Figure 4.**
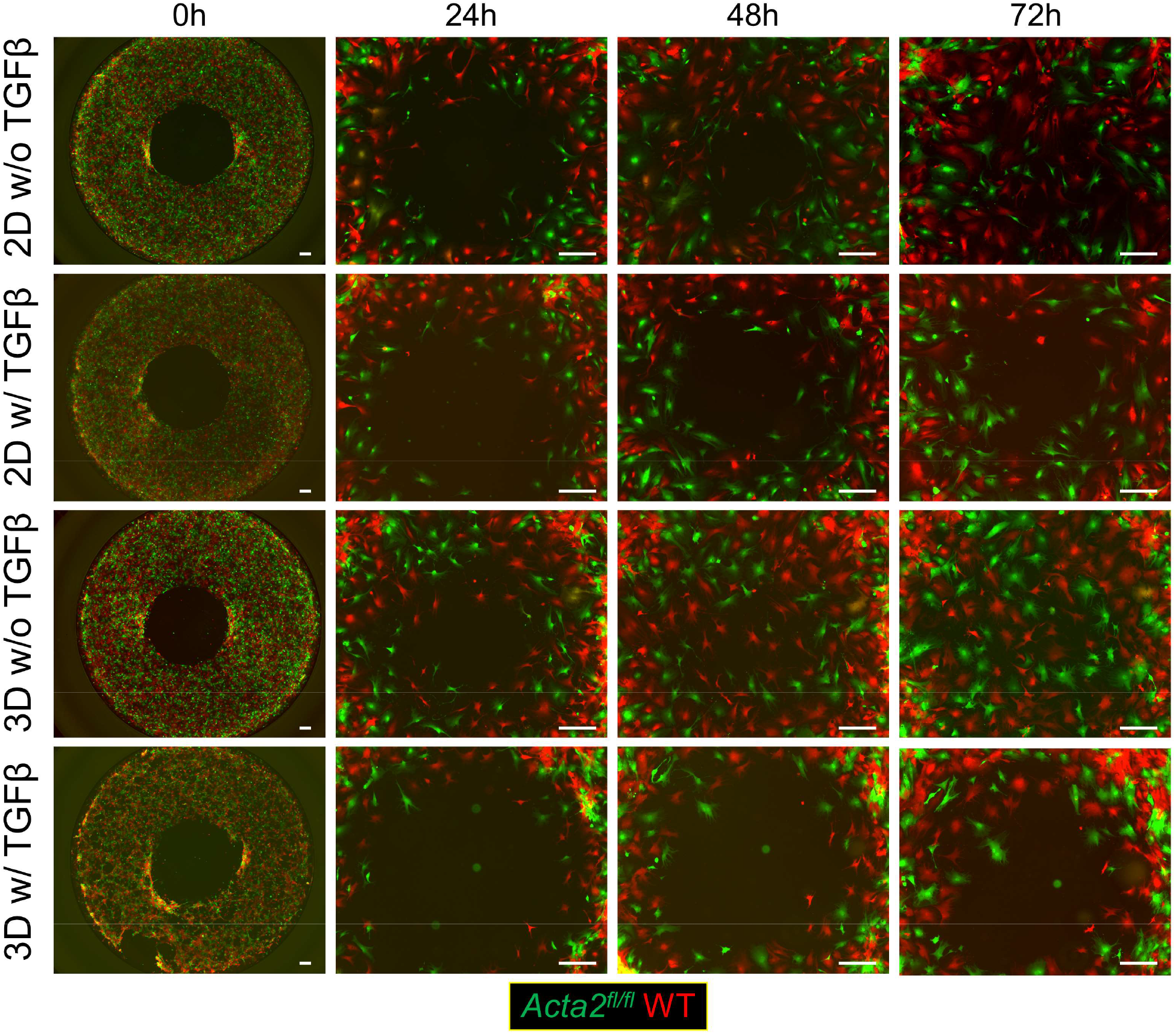
Deletion of *Acta2* does not affect the motility of cardiac fibroblasts. Cardiac fibroblasts isolated from *R26*^*tdTomato*^ (WT) and *Acta2*^*fl/fl*^*;R26*^*eGFP*^ (*Acta2*^*fl/fl*^) mice were transduced with Adeno-Cre and seeded onto the same wells of 96-well plates at a 1:1 ratio with stoppers placed in the center of wells preventing the attachment of cells to the center area of wells. The stoppers were removed after overnight incubation. The migration of cells into the center of wells was monitored every day. For 2D migration, fresh medium with or without TGFβ supplementation was used in the assay. For 3D migration, a layer of collagen gel was overlayed onto the cells after removing the stoppers, followed by the addition of fresh medium with or without TGFβ supplementation. Images represent 3 independent replications. Scale bar: 300 µm.

**Figure 5.**
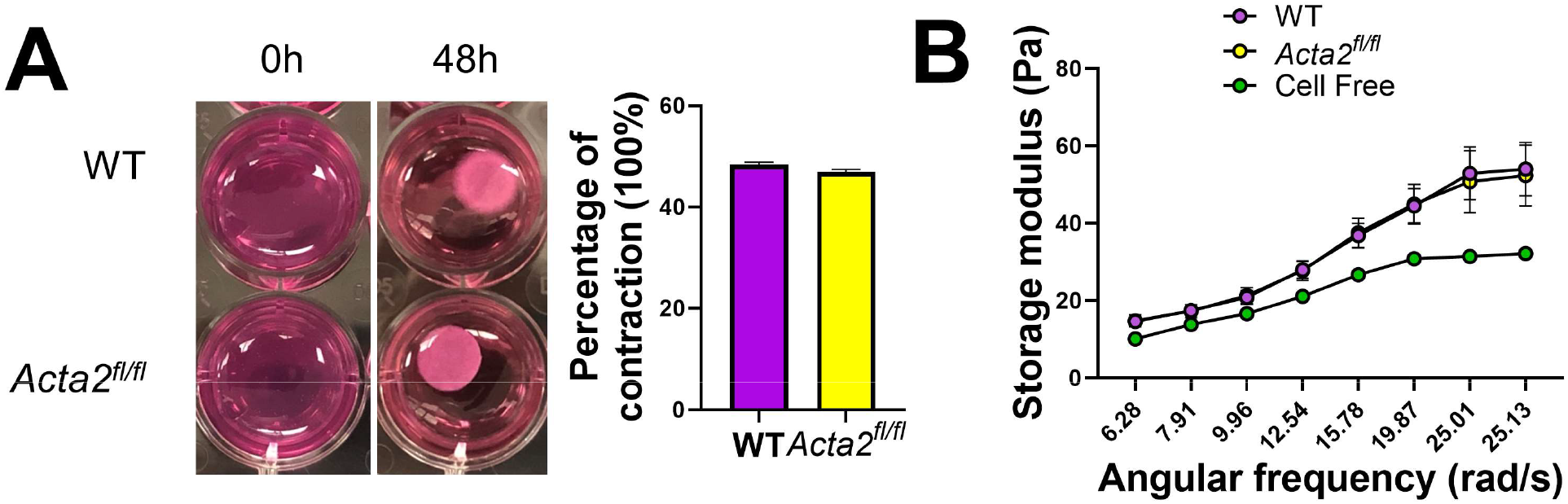
Deletion of *Acta2* does not affect the contractility of cardiac fibroblasts and their ability to stabilize the matrix. Cardiac fibroblasts isolated from WT and *Acta2*^*fl/fl*^ mice were transduced with Adeno-Cre, treated with TGFβ for 2 days, mixed with collagen gel, and poured onto 24 well plates. (A) Gels were released from wells after 12 hours of incubation. Images and quantification show the degree of contraction of gels 48 hours after the release of gels. n=3. (B) The storage moduli of cell-free gels and cell-laden gels 12 hours after incubation were measured using a rheometer. n=5.

### *Acta2* deletion in cardiac fibroblasts does not prevent myofibroblast differentiation

Studies using global *Acta2* KO mice showed that the deletion of *Acta2* did not seem to prevent the formation of F-actin stress fiber in myofibroblasts derived from fibroblasts of several different organs (17-19). To test if this was also the case in cardiac fibroblasts, cardiac fibroblasts were isolated from *Acta2*^*fl/fl*^ and WT mice, treated with Adeno-Cre, and induced for myofibroblast differentiation using TGFβ. Immunocytochemical staining (ICC) using an antibody against SMαA and phallodin staining showed that the amount of F-actin stress fibers was not significantly different between WT and *Acta2-*null cardiac myofibroblasts (**Figure 6A**). To specifically study the effect of *Acta2* deletion on the myofibroblast differentiation of *Tcf21* lineage-traced cardiac fibroblasts, cardiac fibroblasts were isolated from tamoxifen-treated *Tcf21*^*MCM/+*^;*Acta2*^*fl/fl*^;*R26*^*eGFP*^ and *Tcf21*^*MCM/+*^;*R26*^*eGFP*^ mice and cultured in a medium supplemented with TGFβ. ICC showed the lack of SMαA expression in most *Tcf21* lineage-traced cardiac myofibroblasts from *Tcf21*^*MCM/+*^;*Acta2*^*fl/fl*^;*R26*^*eGFP*^ mice but not in *Tcf21* lineage-traced cardiac myofibroblasts from *Tcf21*^*MCM/+*^;*R26*^*eGFP*^ mice (**Figure 6B**). However, no difference was observed in the total F-actin level between *Tcf21* lineage-traced cardiac myofibroblasts of the 2 groups of mice (**Figure 6B**). In the meantime, no difference in the expression of SMαA or F-actin level was observed between non-*Tc21* lineage-traced cardiac myofibroblasts of the 2 groups of mice (**Figure 6B**). Similarly, on day 7 after MI the amount of F-actin stress fibers in *Tcf21* lineage-traced cardiac fibroblasts in the infarct scar of *Tcf21*^*MCM/+*^;*Acta2*^*fl/fl*^;*R26*^*eGFP*^ mice was similar compared to that in *Tcf21*^*MCM/+*^;*R26*^*eGFP*^ mice (**Figure 6C**). Stress fibers guide the maturation of focal adhesion (25). We then questioned if the reduced amount of stress fibers in *Acta2*-null cardiac fibroblasts had an effect on focal adhesion. ICC using an anti-vinculin antibody showed that *Acta2* deletion did not significantly affect the abundance of mature focal adhesion (**Figure 6D**). These results suggest that the deletion of *Acta2* in cardiac fibroblasts does not prevent myofibroblast differentiation.

**Figure 6.**
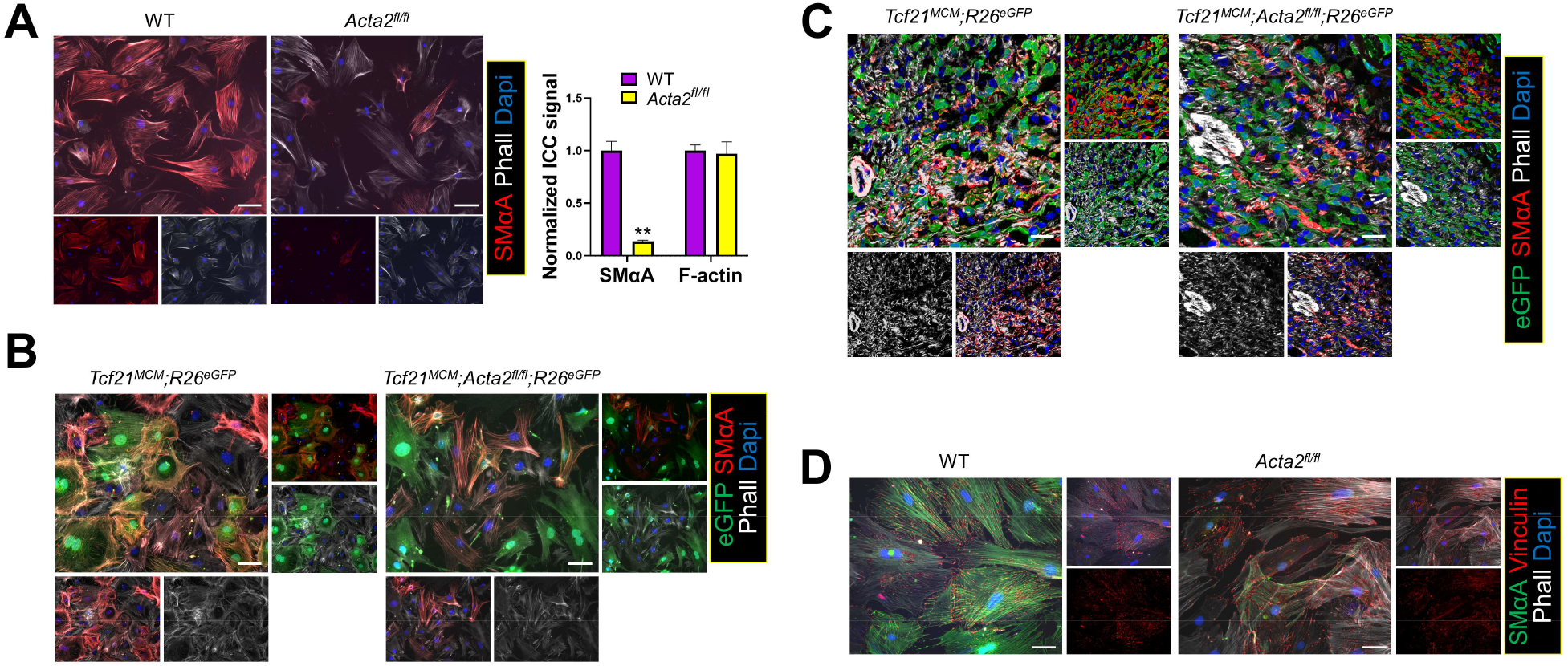
Deletion of *Acta2* does not prevent the myofibroblast differentiation of cardiac fibroblasts. Cardiac fibroblasts isolated from WT and *Acta2*^*fl/fl*^ mice were transduced with Adeno-Cre, treated with TGFβ for 2 days, and subjected to ICC to identify the expression of SMαA and presence of F-actin using an anti-SMαA antibody and phalloidin (Phall), respectively. Nuclei are shown with DAPI. Scale bar: 100 µm. The strength of the signal was determined using ImageJ. n=3. (B) Cardiac fibroblasts isolated from tamoxifen-treated *Tcf21*^*MCM/+*^;*R26*^*eGFP*^ and *Tcf21*^*MCM/+*^;*Acta2*^*fl/fl*^;*R26*^*eGFP*^ mice were treated with TGFβ for 2 days and subjected to ICC to identify the expression of SMαA and presence of F-actin. Nuclei are shown with DAPI. Images represent 3 independent replications. Scale bar: 40 µm. (C) Tamoxifen-treated *Tcf21*^*MCM/+*^;*R26*^*eGFP*^ and *Tcf21*^*MCM/+*^;*Acta2*^*fl/fl*^;*R26*^*eGFP*^ mice were subjected to MI. Heart samples were collected at 7 days post-MI. IHC was performed to identify *Tcf21* lineage-traced (eGFP^+^) cardiac fibroblasts that are positive for SMαA or/and phalloidin. Nuclei are shown with DAPI. Images represent 3 analyzed hearts per group. Scale bar: 20 µm. (D) Cardiac fibroblasts isolated from WT and *Acta2*^*fl/fl*^ mice were transduced with Adeno-Cre, treated with TGFβ for 2 days, and subjected to ICC to identify the expression of SMαA and vinculin using specific antibodies. Nuclei are shown with DAPI. Images represent 3 independent replications. Scale bar: 40 µm. \*\**P* < 0.01.

### Compensatory increase in the expression of non-SMαA actin isoforms in *Acta2-*null cardiac myofibroblasts

Besides *Acta2*/SMαA, 5 other actin isoforms are also expressed in mice, including *Acta1* (skeletal muscle alpha-actin, SkMαA), *Actb* (cytoplasmic beta-actin, CyβA), *Actc1* (cardiac muscle alpha-actin, CMαA), *Actg1* (cytoplasmic gamma-actin, CyγA), and *Actg2* (smooth muscle gamma-actin, SMγA). The normal myofibroblast differentiation of *Acta2*-null cardiac fibroblasts suggests a functional redundancy between different actin isoforms and a possible compensatory increase in the expression of other actin isoforms. To explore these possibilities, we first examined the relative expression levels of different actin isoforms in cardiac fibroblasts after MI using our recently published RNA-seq data obtained using *Tcf21* lineage-traced cardiac fibroblasts isolated from uninjured hearts and infarct scars at different days after MI (26). The result showed that even though *Acta2* was the most upregulated actin isoform in cardiac myofibroblasts after MI, the expression of *Actb, Actg1*, and *Actg2* were also upregulated. Moreover, in cardiac myofibroblasts *Actg1* was expressed at a level comparable to *Acta2* and the expression of *Actb* was twice as high as *Acta2* (**Figure 7A**). Adeno-Cre-and TGFβ-treated WT and *Acta2*^*fl/fl*^ cardiac fibroblasts were then compared to study the impact of *Acta2* on the expression of other actin isoforms by realtime PCR. A unique upregulation in the expression of *Actg2* but not other actin isoforms was identified in *Acta2*-null cardiac myofibroblasts as compared to the WT control (**Figure 7B**). To understand the effect of *Acta2* deletion on the protein level of other actin isoforms, we performed ICC using antibodies against different actin isoforms. ICC using an antibody that recognizes sarcomeric actin isoforms (SkMαA and CMαA) showed that the expression of SkMαA and CMαA in most WT and *Acta2*-null cardiac myofibroblasts was very low (**Figure 7C**), which was consistent with RNA-seq data. The combined protein level of SkMαA and CMαA in *Acta2*-null cardiac myofibroblasts was not significantly different from that in WT cardiac myofibroblasts except for a very small number of *Acta2*-null cardiac myofibroblasts showing an elevated level of SkMαA or/and CMαA, which suggests that sarcomeric actin isoforms do not significantly compensate for the loss of SMαA (**Figure 7C**). Co-staining with an antibody recognizing smooth muscle actin isoforms (SMαA and SMγA) showed that the combined protein level of SMαA and SMγA in *Acta2*-null cardiac myofibroblasts was significantly lower than that in WT cardiac myofibroblasts, suggesting that SMαA is the dominant smooth muscle actin isoform in cardiac myofibroblasts and the increased expression of *Actg2* identified by realtime PCR does not fully compensate for the loss of SMαA (**Figure 7C**). Interestingly, western blot and ICC analyses of Adeno-Cre-and TGFβ-treated WT and *Acta2*^*fl/fl*^ cardiac fibroblasts showed that the total actin protein level was not different between the 2 groups (**Figures 7DE**). Given the limited compensatory increase in the expression of other muscle actin isoforms in *Acta2*-null cardiac myofibroblasts, the unaffected total actin protein level in these cells likely also involved an increased abundance of cytoplasmic actin isoforms, which was possibly due to increased translational efficiency or protein stability of these actin isoforms since no increase in their mRNA expression was identified.

**Figure 7.**
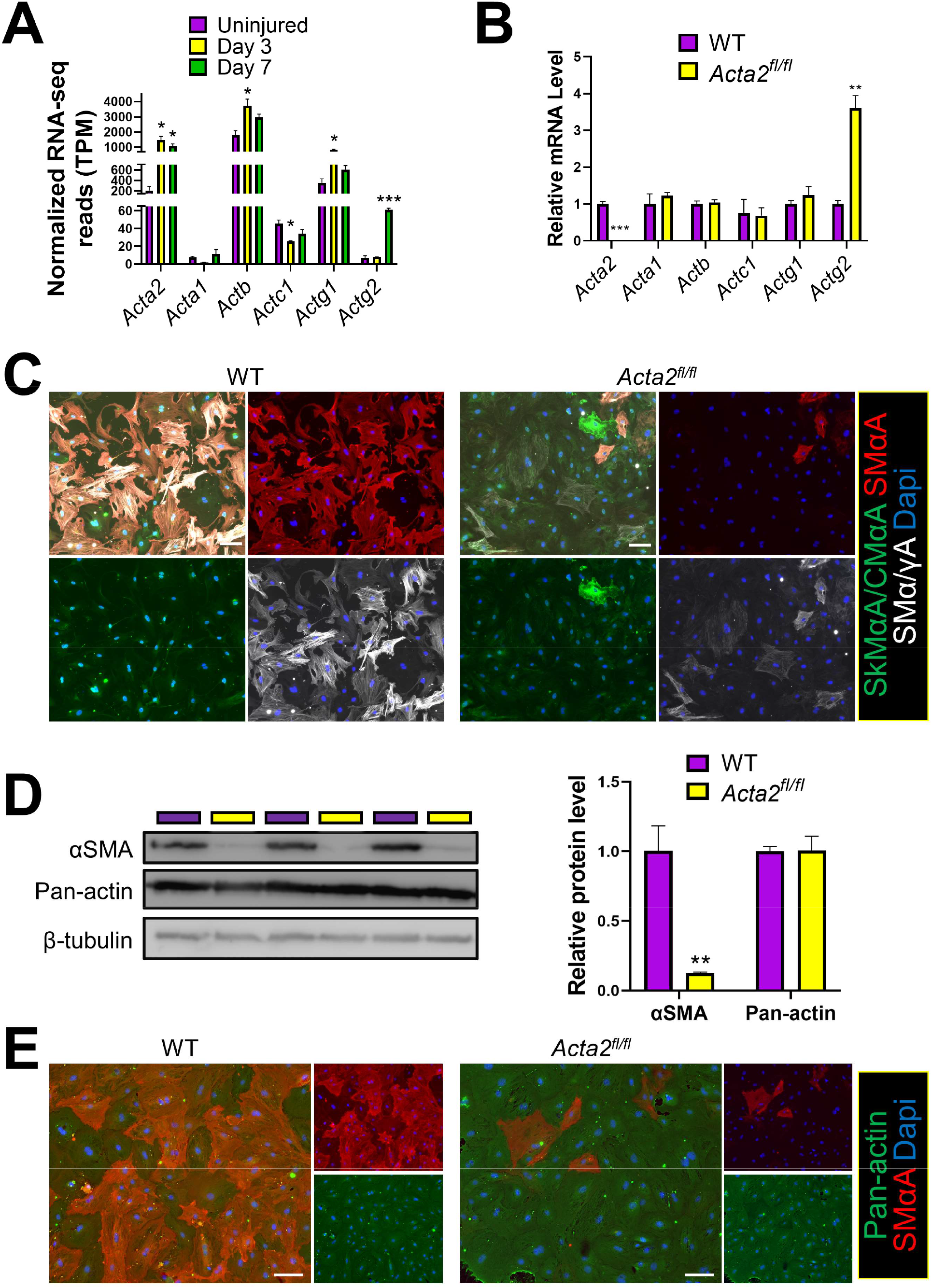
Deletion of *Acta2* leads to a compensatory increase in the expression of other actin isoforms and does not affect the total actin expression level. (A) *Tcf21* lineage-traced cardiac fibroblasts sorted from uninjured hearts and the infarct region at 3 days and 7 days after MI were subjected to RNA-seq. The normalized transcription levels (transcript per million, TPM) of *Acta2, Acta1, Actb, Actc1, Actg1*, and *Actg2* are shown. (B) Cardiac fibroblasts isolated from WT and *Acta2*^*fl/fl*^ mice were transduced with Adeno-Cre and treated with TGFβ for 2 days. RNA was extracted from these cells and used for cDNA synthesis. The transcription levels of *Acta2, Acta1, Actb, Actc1, Actg1*, and *Actg2* were revealed by realtime PCR. (C) Cardiac fibroblasts isolated from WT and *Acta2*^*fl/fl*^ mice were transduced with Adeno-Cre, treated with TGFβ for 2 days, and subjected to ICC to identify the protein level of SMαA, the combined protein level of SkMαA and CMαA, and the combined protein level of SMαA and SMγA. Nuclei are shown with DAPI. Images represent 3 independent replications. Scale bar: 100 µm. (D-E) Cardiac fibroblasts isolated from WT and *Acta2*^*fl/fl*^ mice were transduced with Adeno-Cre and treated with TGFβ for 2 days. Western blot (D) and ICC (E) were performed to quantify SMαA and total actin protein levels using an anti-SMαA antibody and an antibody that recognizes all actin isoforms, respectively. n=3. \**P* < 0.05; \*\**P* < 0.01; \*\*\**P* < 0.0001.

## DISCUSSION

Myofibroblasts with a highly developed cytoplasmic fibrillar system were first identified in skin wounds (27). Later studies found that these cells express an elevated level of SMαA which is incorporated into stress fibers (28-30). Ever since SMαA stress fiber has been used as a marker for myofibroblast differentiation. Due to the specific expression of SMαA in myofibroblasts, it was speculated that SMαA stress fiber may play an important role in multiple myofibroblast activities involved in wound healing such as contraction, motility, and myofibroblast proliferation. However, in this study, using a newly generated mouse line with tamoxifen-inducible cardiac fibroblast-specific deletion of *Acta2*, we identified that the loss of *Acta2* did not significantly affect the myofibroblast differentiation of cardiac fibroblasts. No difference in contractility, matrix stabilization ability, proliferation, or motility was observed between WT and *Acta2-*null cardiac myofibroblasts. In line with results obtained *in vitro*, mice lacking *Acta2* in cardiac fibroblasts also had normal post-MI survival rate, cardiac repair, and cardiac function.

Due to the contractile function of vascular smooth muscle cells, which also express a high level of SMαA, the role of SMαA in the contractility of myofibroblasts received special attention. It was found that overexpression of *Acta2* increased the contractility of 3T3 fibroblasts more significantly than the overexpression of *Actb, Actg1*, or *Actc1*, even though the incorporation of all actin isoforms into stress fibers was identified (31). Disruption of the SMαA stress fiber structure using an SMαA fusion peptide containing the N-terminal of SMαA (32) reduced adhesion and contractility of rat embryo fibroblast cell line REF-52 (15). Besides these early studies of the function of SMαA stress fibers that were mainly performed *in vitro* using cell lines, the function of SMαA stress fibers in primary myofibroblasts and their role during tissue healing were recently studied by a few groups using whole-body *Acta2* KO mice (17-19). Deletion of *Acta2* was reported to reduce the contractility of hepatic stellate cells-derived myofibroblasts (19). However, it was found that mice lacking *Acta2* had normal skin wound closure and the contractility of skin fibroblasts isolated from these mice was not different from those isolated from WT mice (18), which was similar to the effect of *Acta2* KO on cardiac fibroblasts observed in the current study.

Besides the contractile function of myofibroblasts, previous studies have also investigated the effects of SMαA stress fibers on myofibroblast motility, proliferation, and ECM production. A study inhibiting SMαA function using a neutralization antibody identified that the motility of breast tissue-derived myofibroblasts was enhanced when SMαA function was inhibited (14). Another study found that renal fibroblasts isolated from *Acta2*-null mice had enhanced motility and proliferation *in vitro* (17). In addition, the same study also identified an elevated level of collagen production by *Acta2*-null renal myofibroblasts, which exacerbated renal fibrosis. In contrast, the deletion of *Acta2* reduced the collagen expression by myofibroblasts *Acta2*-null hepatic stellate cells (19). Our study, however, did not identify a significant change in any of these myofibroblast functions and activities in cardiac myofibroblasts owing to *Acta2* deletion. Moreover, a study reported that cells increased the stability of the matrix through acting as a crosslinker (20). Stress fibers play an important role in the maturation of focal adhesion, the contact point between cells and matrix (14, 25). The actin stress fiber network in cardiac myofibroblasts is interconnected with the surrounding ECM through focal adhesion and likely function as a structural support lattice to stabilize the early infarct scar before the scar ECM is mature enough to prevent cardiac rupture. The lack of difference in the abundance of stress fiber between WT and of *Acta2* KO cardiac fibroblasts likely also explains the normal survival rate of cardiac fibroblast-specific *Acta2* KO mice.

The reason for the large discrepancy between results generated by different studies of the SMαA function may be multifaceted. First, it is well known that immortalized cell lines, which were used by many early studies, often act significantly differently from primary cells. Second, the experimental procedure of studies using SMαA fusion peptide and neutralization antibodies often included acute treatment of myofibroblasts with SMαA fusion peptide or antibody to disrupt SMαA stress fibers followed by functional analyses. This strategy likely did not provide other actin isoforms with enough time to develop a compensatory effect. Indeed, even though *Acta2*-null mice had normal skin wound contraction (18), a study applying SMαA fusion peptide to skin wounds *in vivo* significantly inhibited rat skin wound contraction (33). Third, the inconsistent results of research focusing on fibroblasts or fibroblast-like cells in different organs using *Acta2* KO mice likely reflect the difference in the adaptation of these cells to *Acta2* deletion. Unlike *Acta2*-null cardiac and skin myofibroblasts that have normal contractility, a study using *Acta2* KO mice reported a reduction in the contractility of myofibroblasts derived from hepatic stellate cells (19). In the current study, a 3-fold increase in the transcription of *Actg2*, the only other actin isoform expressed in smooth muscle, was identified in *Acta2*-null cardiac myofibroblasts as compared to WT cardiac myofibroblasts. A similar increase in the expression of other muscle actin isoforms was also identified in *Acta2*-null skin myofibroblasts when compared to the WT control (18). A similar increase in the expression of non-SMαA muscle actin isoforms was however not identified in myofibroblasts derived from *Acta2*-null hepatic stellate cells (19). Instead, an increase in the expression of cytoplasmic actin isoforms was detected in these cells compared to their WT counterparts. It is possible that without the guidance of stress fibers formed by muscle actin isoforms the ability of non-muscle actin isoforms to form stress fibers is relatively limited, which likely contributed to the more impacted contractile function of *Acta2*-null hepatic stellate cell-derived myofibroblasts.

It is worth noting that even though the deletion of *Acta2* in cardiac fibroblasts does not prevent their myofibroblast differentiation, it does mean that actin stress fibers or the myofibroblast state are dispensable for the post-MI cardiac repair. In contrast, due to the importance of myofibroblasts, a sophisticated gene expression network is likely present in cardiac fibroblasts to ensure their successful myofibroblast differentiation even when the expression of certain myofibroblast functional genes is disrupted. A study disrupting the formation of stress fibers by all actin isoforms is needed in the future to better decipher the function of actin stress fibers in myofibroblasts and post-injury tissue repair.

Taken together, our results indicate that the expression of *Acta2*/SMαA is not required for the myofibroblast differentiation of cardiac fibroblasts and their activities in post-MI cardiac repair, which is at least partially due to the compensatory increase in the expression of other actin isoforms and the functional redundancy between SMαA and non-SMαA actin isoforms. Our study also suggests that significant heterogeneity is present within the over-simplified fibroblast and fibroblast-like cell types.

## Method

### Mice

Mouse embryonic stem cells with a “knockout-first” *Acta2* allele purchased from European Mouse Mutant Cell Repository were used to generate chimeric mice. Chimeric mice were crossed with WT C57BL/6 mice to obtain offsprings carrying the mutant allele which were then crossed with mice expressing FLPe recombinase (Jackson Laboratory, #003946) to delete the LacZ-neo cassette upstream of exon 5 of *Acta2* to generate *Acta2*^*fll+*^ mice with exons 5 to 7 of *Acta2* flanked by *loxP* sites. *Tcf21*^*MCM/+*^ (34) and *R26*^*eGFP*^ (35) mice were crossed to generate *Tcf21*^*MCM/+*^;*R26*^*eGFP*^ mice. *Acta2*^*fll+*^ mice were crossed with *Tcf21*^*MCM/+*^;*R26*^*eGFP*^ mice to generate *Tcf21*^*MCM/+*^;*Acta2*^*fl/fl*^;*R26*^*eGFP*^ mice. *R26*^*tdTomato*^ mice were purchased from Jackson Laboratory (#007914).

### Antibodies and biologics

Anti-eGFP (ab13970) and anti-Col1(ab34710) antibodies were purchased from Abcam. Anti-Ki67 antibody (9129S) and Alexa Fluor® 647 Phalloidin (8940S) were purchased from Cell Signaling Technology. Anti-SMαA (113200), anti-SkMαA/CMαA (A2172), and anti-SMα/γA (A7607) antibodies were purchased from Millipore Sigma. Pan anti-actin and anti-β-tubulin antibodies were purchased from DSHB. Anti-vinculin (NB600-1293) antibody was purchased from Novus. Goat anti-mouse IgG2a Alexa Fluor 488 (A-21131), goat anti-mouse IgG2b Alexa Fluor 488 (A-21141), goat anti-mouse IgM Alexa Fluor 488 (A-21042), goat anti-mouse IgG2a Alexa Fluor 555 (A-21137), goat anti-mouse IgG1 Alexa Fluor 555 (A-21127), goat anti-rabbit IgG Alexa Fluor 555 (A-21428), goat anti-mouse IgG Alexa Fluor 647 (A-21235), and goat anti-chicken IgY Alexa Fluor 488 (A-11039) antibodies were purchased from Thermo Scientific. EdU (catalog sc-284628A) was purchased from Santa Cruz Biotechnology Inc. Collagenase D (catalog 11088866001) and Dispase II (catalog 4942078001) were purchased from Roche Diagnostics. Collagen, type I (356236) was purchased from Corning. TGF-β (240-B) was purchased from R&D Systems.

### Animal procedures

To induce the activity of the MerCreMer protein, mice were treated with tamoxifen (MilliporeSigma, T5648) dissolved in corn oil through i.p. injections or gavage at a dosage of 75 mg/kg body weight/day for 5 days starting at 6 weeks of age. Mice were subjected to permanent surgical ligation of the left coronary artery to induce MI at 9 weeks of age (36). Briefly, mice were anesthetized using isoflurane and a left lateral thoracotomy was performed. The left coronary artery was identified and ligated just below the left atrium. To ensure the efficient deletion of *Acta2*, mice were given another course of tamoxifen treatment from 1 day before MI to 5 days after MI. Echocardiography was performed in M-mode using a Toshiba Aplio SSA-770a ultrasound system and a 12-MHz transducer as previously described (37). For pain management related to surgical procedures, mice were given a dose of Carprofen (5 mg/kg body weight) before the surgery followed by a second dose 12 hours after the surgery. Mice were checked every day after MI for survival rate. Mice found dead were subjected to necropsy to identify cardiac rupture.

### EdU in vivo fibroblast proliferation assay

Mice were treated with EdU at a dosage of 50 mg/kg body weight through i.p. injections. Four hours after EdU injections, mice were sacrificed, and heart samples were collected. EdU detection was carried out after IHC staining using the Click-iT Plus Alexa Fluor 647 Picolyl Azide Toolkit (C10643) from Thermo Fisher Scientific.

### Cell isolation and culture

Cardiac fibroblasts were isolated as previously described (3). Briefly, heart tissue was minced and digested in DMEM containing 0.75 U/ml collagenase D (Roche, 11088866001), 1.0 U/ml Dispase II (Roche, 10165859001), and 1 mM CaCl_2_ at 37°C for 40 minutes. The slurry was then passed through a 100 μm cell strainer and then a 40 μm cell strainer. Cells were collected by centrifugation at 350 *g* for 10 minutes. The cell pellets were resuspended in a growth medium composed of DMEM with 10% bovine growth serum (BGS) and 1% of an antibiotic mixture containing 10,000 U/ml penicillin, 10 mg/ml streptomycin, and 25 μg/ml amphotericin B and seeded on cell culture plates. To induce *Acta2* deletion, WT and *Acta2*^*fl/fl*^ cardiac fibroblasts were treated with Adeno-Cre. Myofibroblast differentiation was induced by TGFβ (10 ng/ml) treatment for 2 days.

### Gel contraction assay

Gel contraction assay was performed as previously described with some modifications. Briefly, 50,000 cardiac fibroblasts that had been treated with TGFβ (10 ng/ml) and Adeno-Cre were resuspended in 0.4 ml of growth medium and mixed with 0.2 ml of collagen solution that contains 0.1% acetic acid. 4 µl of 1 M NaOH was added to neutralize the acetic acid. 0.5 ml of the mixture was added to each well of 24 well plates and allowed to solidify at room temperature for 30 min, followed by the addition of 0.5 ml of growth medium supplemented with TGFβ. 12 hours after the incubation at 37 °C, gels were carefully released from the well to allow the contraction.

### Rheology

Cell-free and cell-laden collagen gels were made and cultured as described above. After 24 hours of incubation, collagen gels were released and trimmed using a 5 mm biopsy punch. Using a TA Discovery HR-2 rheometer and a 5 mm crosshatched plate, the storage moduli of gels were determined by frequency sweeping from 6.28 to 25.13 (rad/s) at 2% strain and 10% compression.

### Cell motility assay

Cell motility assay was performed using Oris Cell Migration Assembly Kit (Platypus Technologies) following the manufacturer’s protocol with some modifications. Briefly, TGFβ (10 ng/ml) and Adeno-Cre-treated *R26*^*tdTomato*^ and *Acta2*^*fl/fl*^*;R26*^*eGFP*^ cardiac fibroblasts were mixed at a 1:1 ratio. A total of 40,000 cells were seeded into each well of 96 well plates with a stopper placed in the center of the well. Cells were cultured in a maintenance medium containing DMEM with 0.5% BGS and 1% of an antibiotic mixture at 37 °C overnight before the removal of stoppers. A quick rinse with PBS was performed, followed by the addition of a fresh maintenance medium with or without TGFβ. For the 3D migration test, after the rinse, 0.5 ml neutralized collagen solution (3 mg/ml) was added to each well and allowed to solidify before adding fresh maintenance medium with or without TGFβ. The migration of cells into the center of the well was monitored.

### Realtime PCR

Realtime PCR was performed as previously described (38). Briefly, RNA was extracted using a Direct-zol RNA Microprep Kit (Zymo). cDNA was synthesized using an iScript cDNA Synthesis Kit (Bio-Rad). Realtime PCR was carried out using a CFX RT-PCR detection system (Bio-Rad) with SsoAdvanced Universal SYBR Green Supermix (Bio-Rad). Relative mRNA content was normalized to 18S rRNA content. The primers used are listed in table 1.

**Table 1.**
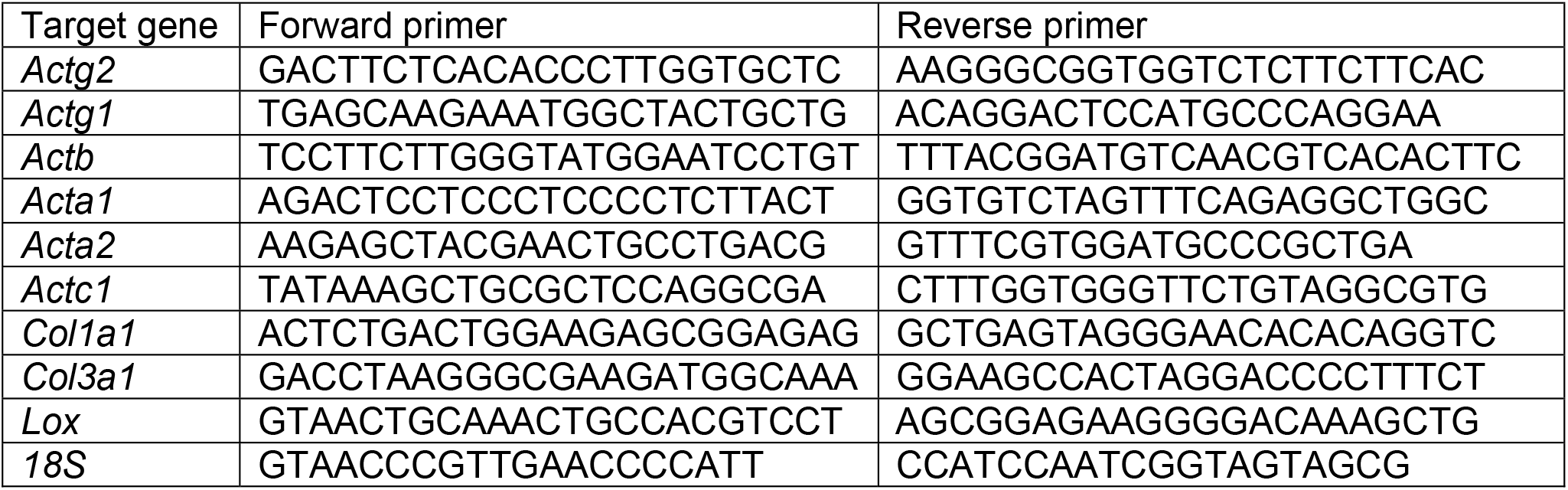
Primer sequences used in realtime PCR.

### Western blot

Western blot was performed as previously described (39) with some modifications. Briefly, cells were lysed in RIPA buffer with Halt Protease Inhibitor Cocktail (Thermo Scientific) to extract the protein. SDS-PAGE and blotting were performed using Bio-Rad Mini-PROTEAN Electrophoresis System. After the immunostaining, images were taken using a ProteinSimple FluorChem R System. Band density was normalized to β-tubulin content.

### Immunocytochemical staining

Cells grown on multiple-chamber slides or multiple well plates were fixed in 4% paraformaldehyde (PFA) for 10 minutes, rinsed 3 times in TBS with 0.1% Triton X-100, incubated in blocking buffer (TBS, 0.1% Triton X-100, and 3% BSA), and then incubated with primary antibodies diluted in blocking buffer overnight at 4°C. Cells were then rinsed in TBS with 0.1% Triton X-100 3 times and stained with corresponding secondary antibodies diluted in blocking buffer for 1 hour at room temperature. Stained slides were then rinsed and mounted in a mounting medium containing DAPI (Vector Laboratories). Images were captured using an Echo Revolve microscope. Signal strength was determined using ImageJ.

### IHC staining

IHC staining was performed as previously described (3). Briefly, mouse heart samples were fixed in 4% PFA, incubated in 30% sucrose dissolved in PBS, and embedded in OCT (Tissue-Tek) for cryosectioning. Cryosections (5 μm thick) were blocked with 5% goat serum and 0.2% Triton X-100 diluted in TBS, incubated in primary antibodies diluted in blocking buffer overnight at 4°C, and then incubated in appropriate fluorophore-conjugated secondary antibodies diluted in blocking buffer for 1 hour at room temperature. Stained sections were then mounted in a mounting medium containing DAPI (Vector Laboratories). Images were captured using a Leica SP8X confocal microscope.

### Trichrome staining

Trichrome staining was performed using reagents for Masson’s Trichrome for Connective Tissue (Electron Microscopy Sciences) following the manufacturer’s protocol.) Percentage of the fibrotic area in the left ventricle and average left ventricle thickness at 3 different segments were determined using ImageJ.

### Statistics

All data are expressed as mean ± SEM unless otherwise stated. Data were analyzed using GraphPad Prism 9 (GraphPad Software, Inc.). Two-tailed *t* test was used to determine the significance of differences between the 2 groups. One-way ANOVA with post hoc Tukey’s test was used to determine the significance of difference when more than 2 groups were compared. Chi-Square Test was used to determine the significance of difference between the survival rates of *Tcf21*^*MCM/+*^;*R26*^*eGFP*^ and *Tcf21*^*MCM/+*^;*Acta2*^*fl/fl*^;*R26*^*eGFP*^ mice. *P* < 0.05 was considered significant.

### Study approval

All experiments involving mice were approved by the IACUC at LSU (approval number IACUC 18-024).

## Author contribution

X.F. conceived the study; Y.L., C.L., Q.L., L.W., and A.B. performed experiments. Y.L., C.L., and J.J.. analyzed data. Y.L, C.L., J.F., J.M., and X.F. interpreted the data, assembled the results, and wrote the manuscript with inputs from all authors.

